# Sex-biased migration and admixture in macaque species revealed by comparison between autosomal and X-chromosomal genomic sequences

**DOI:** 10.1101/2020.05.26.115915

**Authors:** Naoki Osada, Kazunari Matsudaira, Yuzuru Hamada, Suchinda Malaivijitnond

**Author notes:** Correspondence to: Naoki Osada, Kita-14, Nishi-9, Kita-ku, Sapporo, Hokkaido 060-0814, Japan.

## Abstract

The role of sex-specific demography in hybridization and admixture of genetically diverged species and populations is essential to understand the mechanisms forming the genomic diversity of sexually reproducing organisms. In order to infer how sex-linked genetic loci have been differentiated undergoing frequent hybridization and admixture, we examined 17 whole-genome sequences of seven species of the genus *Macaca*, which shows frequent inter-specific hybridization and predominantly female philopatry. We found that hybridization and admixture were prevalent within these species. For three cases of suggested hybrid origin of species/subspecies, *M. arctoides*, *M. fascicularis ssp. aurea*, and Chinese *M. mulatta*, we examined the level of admixture of X chromosomes, which is less affected by male-biased migration than that of autosomes. In one case, we were able to determine that *M. cyclopis* and *M. fuscata* was genetically closer to Chinese *M. mulatta* than to the Indian *M. mulatta*, and the admixture level of Chinese *M. mulatta* and *M. fuscata*/*cyclopis* was more pronounced on the X chromosome than on autosomes. Since the mitochondrial genomes of Chinese *M. mulatta*, *M. cyclopis*, and *M. fuscata* were found to cluster together, and the mitochondrial genome of Indian *M. mulatta* is more distantly related, the observed pattern of genetic differentiation on X-chromosomal loci is consistent with the nuclear swamping hypothesis, in which strong, continuous male-biased introgression from the ancestral Chinese *M. mulatta* population to a population related to *M. fuscata* and *M. cyclopis* generated incongruencies between the genealogies of the mitochondrial and autosomal genomes.

## Introduction

A key issue in evolutionary genetics is understanding the mechanisms by which populations are genetically differentiated and continue diverging to form separate species. Genetic differentiation is often attributed to geographical isolation, and genetically differentiated populations evolve into new, reproductively isolated, species. However, hybridization and admixture among genetically divergent populations occasionally occur, impeding further genetic differentiation. Hybridization and admixture have frequently been observed in populations of wild animals and between different species (Green et al. 2010; Osada et al. 2010; Meyer et al. 2012; Cahill et al. 2015; Fan et al. 2018).

Individuals of different sexes have shown different dispersal patterns in a wide range of sexually reproducing species. Sex-biased migration affects the pattern of genetic differentiation across genomes. In animals with an X-Y sex chromosome system, the X and Y chromosomes, as well as the mitochondrial genomes, can be used as markers to track sex-biased migration. The effect of migration on mitochondrial genomes depends solely upon female migration, while that of Y chromosomes depends solely upon male migration. Similarly, the migration of males is expected to have less impact on X chromosomes than on autosomes, because a female carries two X chromosomes but a male has only one. Genetic differentiation at these sex-linked markers, therefore, provides insights into the way in which sex-biased migration has shaped animal genomes through hybridization and admixture among populations and species (Heyer and Segurel 2010).

In order to investigate patterns of genetic differentiation across genomes during hybridization process, and to estimate how sex-biased migration contributes to species divergence, we focused on macaque monkeys. The genus *Macaca* consists of 24 species distributed in the part of the Eurasian and African continents, mostly in the South, Southeast, and East Asia (Zinner 2013; Roos et al. 2014). They are classified into four to seven species groups, depending on the criteria used (Zinner 2013; Roos et al. 2019). In this study, we followed the grouping proposed by Zinner et al. (2013), and focused on four out of seven species groups: the *fascicularis* group, including *Macaca fascicularis* (the cynomolgus or long-tailed macaque); the *mulatta* group including *M. mulatta* (the rhesus macaque), *M. cyclopis* (the Taiwanese macaque), and *M. fuscata* (the Japanese macaque); the *sinica* group, including *M. sinica* (the toque macaque), *M. thibetana* (the Tibetan macaque), *M. assamensis* (the Assamese macaque), and *M. radiata* (the bonnet macaque); and the *arctoides* group, including *M. arctoides* (the stump-tailed macaque). Genome-wide phylogenetic analysis showed that the *fascicularis* and *mulatta* groups and the *sinica* and *arctoides* groups are sister pairs (Fan et al. 2018).

Previous studies have revealed that ancient inter-specific gene flow has been common in macaques, not only between sister species such as *M. fascicularis* and *M. mulatta* (Osada et al. 2010; Yan et al. 2011) but also between different species groups, for example, *M. thibetana* in the *sinica* group and *M. mulatta* in the *mulatta* group (Fan et al. 2014). Ancient hybridization between the *arctoides* and *mulatta* groups was also supported by whole-genome sequencing analysis of *M. arctoides* (Fan et al. 2018). The divergence of the *sinica-arctoides* and *fascicularis-mulatta* groups is estimated to have occurred around 1.5–2.0 Mya, as per fossil evidence (Delson 1980), but much earlier estimates (2.8–4.9 Mya) have been obtained using nucleotide sequences (Perelman et al. 2011; Jiang et al. 2016; Matsudaira et al. 2018; Roos et al. 2019).

In some macaque species, discordant genealogies have been obtained using autosomal, mitochondrial, and Y-chromosomal loci, a situation potentially reflecting sex-biased migration patterns in the past (Tosi, Morales, and Melnick 2000; Tosi, Morales, and Melnick 2002; Tosi, Morales, and Melnick 2003; Evans et al. 2010; Zinner, Arnold, and Roos 2011). In our focal four species groups, three cases of incongruence between mitochondrial genealogies and the conventional taxonomic classification have been shown. (1) The mitochondrial genome of *M. arctoides* was a sister to the *mulatta*-group species in the mitochondrial phylogenetic tree (Fan et al. 2018; Roos et al. 2019); however, a genome sequencing study of *M. arctoides* showed that the *M. arctoides* genome had a strong affinity to the genomes of *sinica*-group species (Fan et al. 2018). (2) *M. fascicularis* ssp. *aurea* has been found to be distributed in Myanmar and Thailand, having a morphology distinct from the other nine subspecies (Fooden, 1995). A genetic study using mitochondrial and Y-chromosomal sequences showed that *M. fascicularis* ssp. *aurea* clustered with the *sinica* group at the mitochondrial locus, but clustered with the *fascicularis* group at the Y-chromosomal locus (Matsudaira et al. 2018). (3) The mitochondrial sequences of *M. fuscata* and *M. cyclopis* are determined to be more similar to the Chinese *M. mulatta* than to the Indian *M. mulatta*, contradicting the conventional taxonomic classification of *M. mulatta* as a single species (Melnick et al. 1993; Matsudaira et al. 2018; Roos et al. 2019).

Incongruent phylogenies constructed using sex-linked loci have been often considered to be the result of male-biased migration in macaques. Because current macaque populations show predominantly female philopatry, the lineage of mitochondria is assumed to reflect the genetic lineage of the recipient population of migrants, whereas the other autosomal and Y-chromosomal genomes in the recipient population may have been “swamped” by the genomes of donor (introgressing) populations (Zinner, Arnold, and Roos 2011; Zinner et al. 2013). This is referred to as the nuclear swamping hypothesis. A schematic representation of the nuclear swamping hypothesis is presented in Figure 1. However, in many species other than primates, introgression of mitochondrial genomes between species, so-called mitochondrial capture, is not rare (Bachtrog et al. 2006; Veale, Russell, and King 2018). Since the effective population size of the mitochondrial genome is much smaller than that of autosomal genomes, introgressed mitochondrial genomes might easily become fixed in the recipient population by genetic drift (Osada 2011). Besides, in several animal species, paternal leakage of mitochondrial genomes has been observed, particularly in cases of inter-specific hybridization (Rokas, Ladoukakis, and Zouros 2003; Mastrantonio, Urbanelli, and Porretta 2019). Therefore, in order to determine the factors underlying incongruent genealogies between autosomal and sex-linked loci, a study focusing on sex-biased migration using loci other than mitochondrial locus is desirable. Genetic differentiation at the X chromosome has the potential to provide a picture with substantially higher resolution in the estimation of sex-biased migration rates, because X chromosomes are longer than mitochondria and, more importantly, recombine except at the pseudo-autosomal regions (Figure 1) (Osada et al. 2013; Evans et al. 2017; Goldberg et al. 2017).

**Figure 1.**
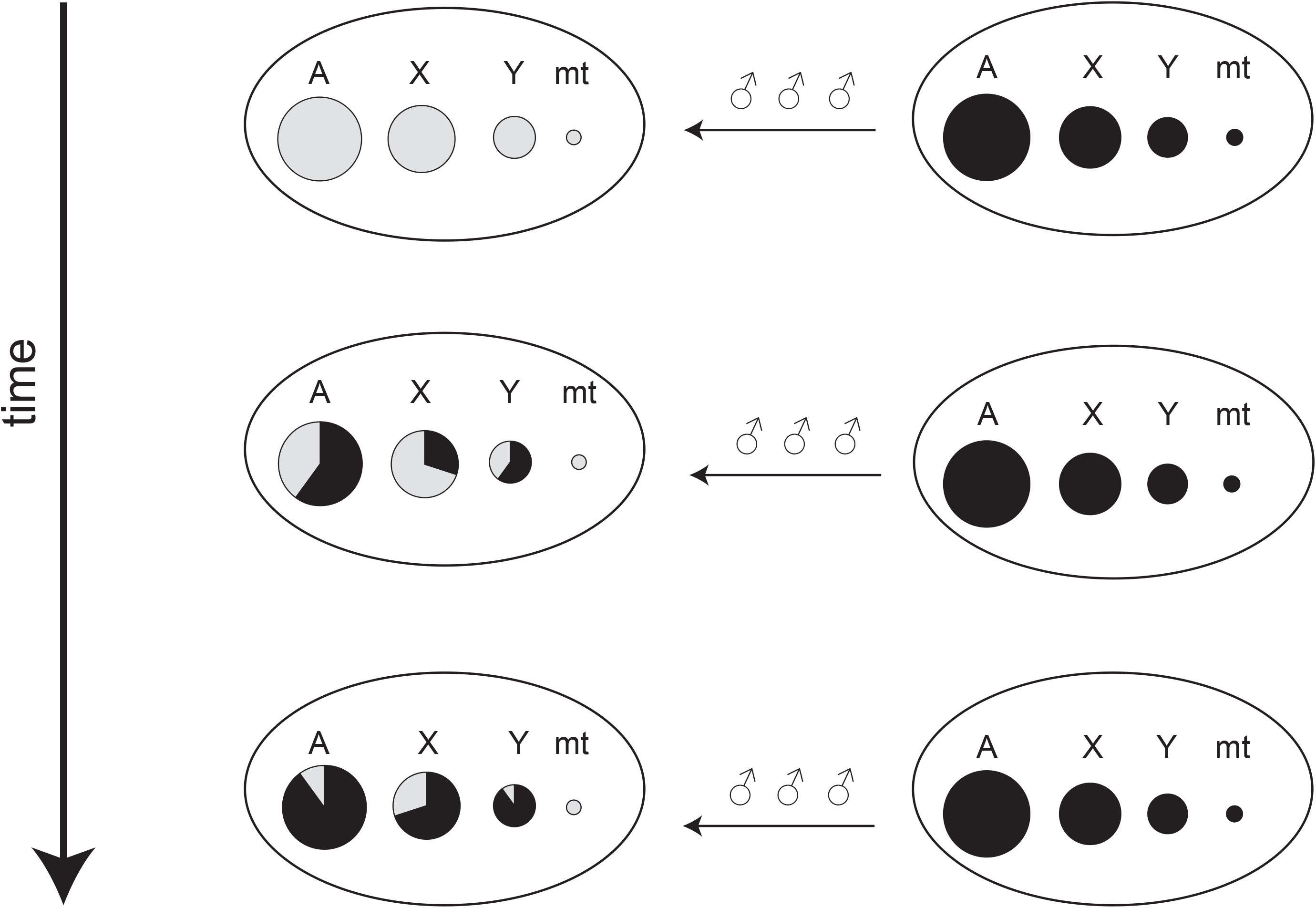
Schematic representation of the nuclear swamping hypothesis. The letters A, X, Y, and mt represent autosomal, X-chromosomal, Y-chromosomal, and mitochondrial genomes, respectively, and the circles indicate the gene pool of a population. When strong male-biased migration continues for a long time, the nuclear genomes of the recipient population are replaced by the donor alleles, while mitochondrial genomes retain the original (gray) genetic components. However, if the replacement is not complete, we would expect that the X chromosome would retain more original genetic components than the autosomes.

In order to investigate how sex-biased migration affects the inter-specific hybridization of genomes, we used the above-mentioned three cases that showed incongruent species trees between mitochondrial and nuclear loci: (1) between *M. arctoides* and the *mulatta-*group species; (2) between *M. fascicularis* ssp. *aurea* and the *sinica*-group species; and (3) among the *mulatta*-group species. We clarified the phylogeny of eight macaque species using 17 whole-genome sequences, by combining newly and previously determined whole-genome sequences of macaques, and compared the level of genetic admixture at the mitochondrial genome, autosomes, and X chromosomes.

## Results

### Whole-genome sequencing and genetic diversity in populations

This study has performed whole-genome sequencing of five macaque samples: one female *M. fascicularis* ssp. *aurea* (CMA1), one male *M. fascicularis* ssp. *aurea* (CMA2), one male *M. fascicularis* ssp. *fascicularis* from Southern Thailand (CMT1), one male *M. fuscata* (JPM1), and one male *M. cyclopis* (TWM1) (Supplementary Table 1). We also obtained short-read sequences of an additional 12 samples from previous studies (Supplementary Table 2): three Chinese *M. mulatta* (RMC1–3), two Indian *M. mulatta* (RMI1–2), three Mauritian *M. fascicularis* (CMM1–3), one Vietnamese *M. fascicularis* (CMV1), one Chinese *M. assamensis* ssp. *assamensis* (ASM1), one *M. thibetana* (TIM1), and one *M. arctoides* from China (STM1). CM and RM represent cynomolgus and rhesus macaques, respectively. The abbreviations of the sample names are summarized in Table 1. These symbols are used to represent the names of the samples.

**Table 1.** Summary of whole-genome sequencing

| sample | symbol | coverage | heterozygosity | reference |
| --- | --- | --- | --- | --- |
| *M. fascicularis* (Thailand) | CMT1 | 39.8 | 0.0034 | This study |
| *M. fascicularis* (Vietnam) | CMV1 | 32.0 | 0.0029 | Yan et al. 2011 |
| *M. fascicularis* (Mauritius) | CMM1 | 36.8 | 0.0025 | Eliscen et al. 2014 |
| *M. fascicularis* (Mauritius) | CMM2 | 35.5 | 0.0024 | Eliscen et al. 2014 |
| *M. fascicularis* (Mauritius) | CMM3 | 39.6 | 0.0026 | Eliscen et al. 2014 |
| *M. fascicularis* ssp. *aurea* | CMA1 | 45.7 | 0.0013 | This study |
| *M. fascicularis* ssp. *aurea* | CMA2 | 41.7 | 0.0029 | This study |
| *M. mulatta* (China) | RMC1 | 34.4 | 0.0025 | Xue et al. 2016 |
| *M. mulatta* (China) | RMC2 | 23.5 | 0.0025 | Xue et al. 2016 |
| *M. mulatta* (China) | RMC3 | 29.3 | 0.0024 | Xue et al. 2016 |
| *M. mulatta* (India) | RMI1 | 40.2 | 0.0023 | Xue et al. 2016 |
| *M. mulatta* (India) | RMI2 | 47.6 | 0.0024 | Xue et al. 2016 |
| *M. fuscata* | JPM1 | 27.5 | 0.0010 | This study |
| *M. cyclopis* | TWM1 | 40.0 | 0.0016 | This study |
| *M. thibetana* | TIM1 | 38.1 | 0.0009 | Fan et al. 2014 |
| *M. assamensis* | ASM1 | 50.8 | 0.0029 | Fan et al. 2018 |
| *M. arctoides* | STM1 | 33.2 | 0.0017 | Fan et al. 2018 |

Because we focused on admixture between different species groups, the reads were mapped to the reference genome sequence of the olive baboon (*Papio anubis*, OLB), which should be equally distant from all the analyzed samples, in order to avoid mapping bias due to short reads. The mapping rate to the OLB genome was greater than 99 % except for one sample, RMI2 (Supplementary Figure 1).

The coverage of the genomes and the observed heterozygosity for each sample are summarized in Table 1. CMA1, JPM1, and TIM1 have showed very low levels of heterozygosity (0.0009–0.0013), while CMA2, CMT1, CMV1, and ASM1 were determined to have the highest levels of heterozygosity (0.0029–0.0034). In order to evaluate the low observed genetic diversity of CMA1, JPM1, and TIM1, we calculated the number of run of homozygosity (ROH) regions greater than 500kb length for each sample (Supplementary Table 3). Although the heterozygosity of CMA1 was higher than that of JPM1, the number of ROH regions in CMA1 was noticeably higher than that in JPM1 (195 vs. 13), suggesting that the low heterozygosity of CMA1 may be due to a recent inbreeding effect.

### Species phylogeny

To examine the phylogenetic relationships among the macaques, we first constructed a tree using mitochondrial genomes (Figure 2). Complete mitochondrial genome sequences were assembled for the five newly sequenced samples (CMT1, JPM1, TWM1, CMA1, and CMA2) and four additional samples (CMV1, ASM1, STM1, and TIM1). Additional 31 mitochondrial genomes of macaques were directly downloaded from the public database or assembled using short-read sequences in the public database (Supplementary Table 4). As shown in Figure 2, at the mitochondrial locus, *M. arctoides* clustered with the *mulatta* group, while *M. fascicularis* ssp. *aurea* clustered with the *sinica* group. The tree also indicated that *M. mulatta* samples were paraphyletic; the Indian *M. mulatta* samples were considered an outgroup for the Chinese *M. mulatta*, *M. fuscata*, and *M. cyclopis* samples. The result of our study is largely consistent with the result of the previous ones (Roos et al. 2019).

**Figure 2.**
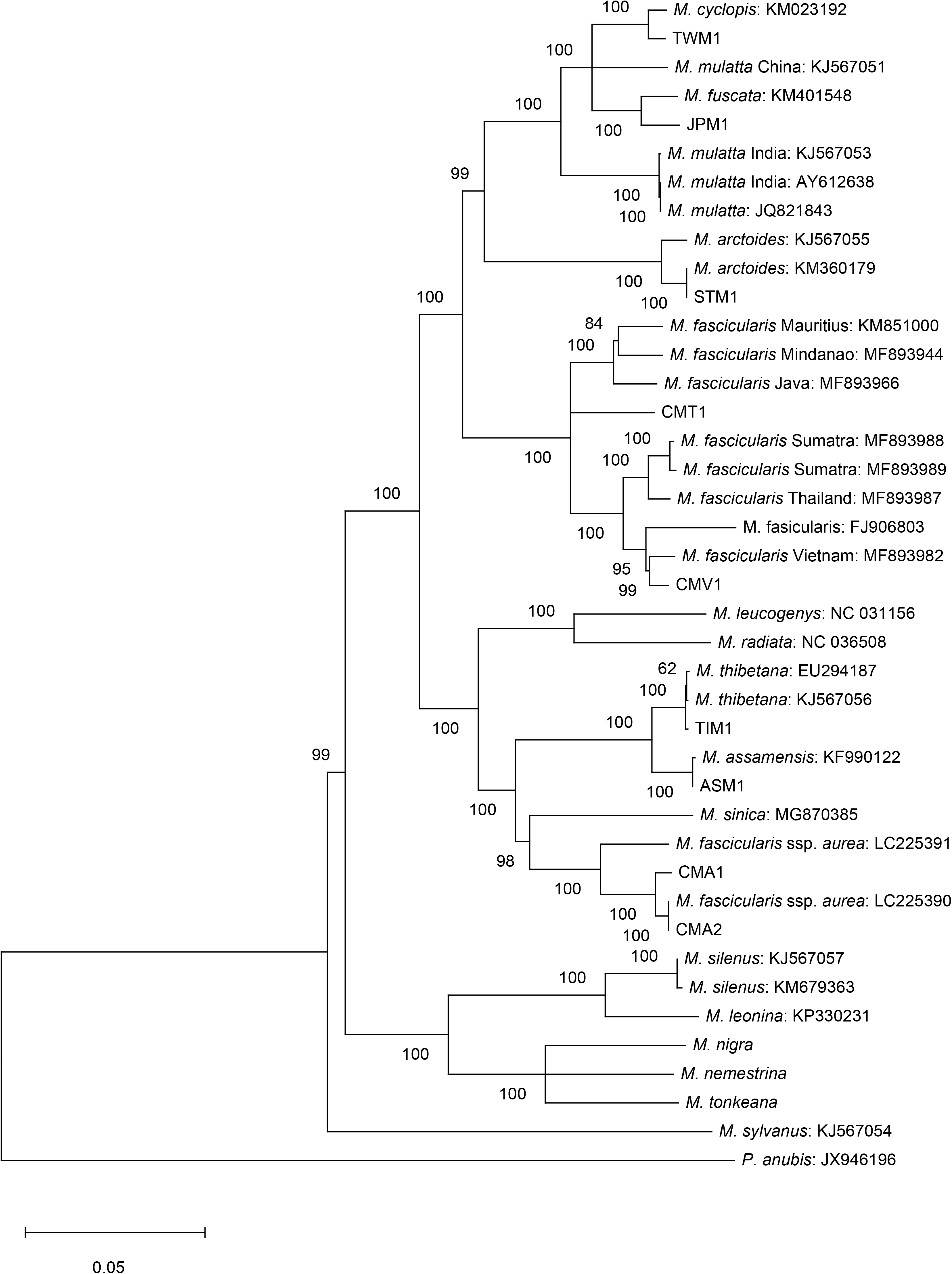
Mitochondrial genealogy, including publicly available mitochondrial genome sequences. Bootstrap percentile values are shown near the nodes.

Next, we constructed a neighbor-joining tree using the genetic distance between nuclear genomes with the draft genome sequence of OLB as an outgroup (Figure 3). The tree showed the sister relationship between the *sinica* and *arctoides* groups, and between the *fascicularis* and *mulatta* groups, as presented in previous studies (Perelman et al. 2011; Fan et al. 2014; Fan et al. 2018). We also reconstructed a tree using the TreeMix software (Pickrell and Pritchard 2012) and a graph using the neighbor-net method (Bryant and Moulton 2004). These results were found to be essentially the same as the neighbor-joining tree (Supplementary Figures 2 and 3) but showed a signature of gene flow between species.

**Figure 3.**
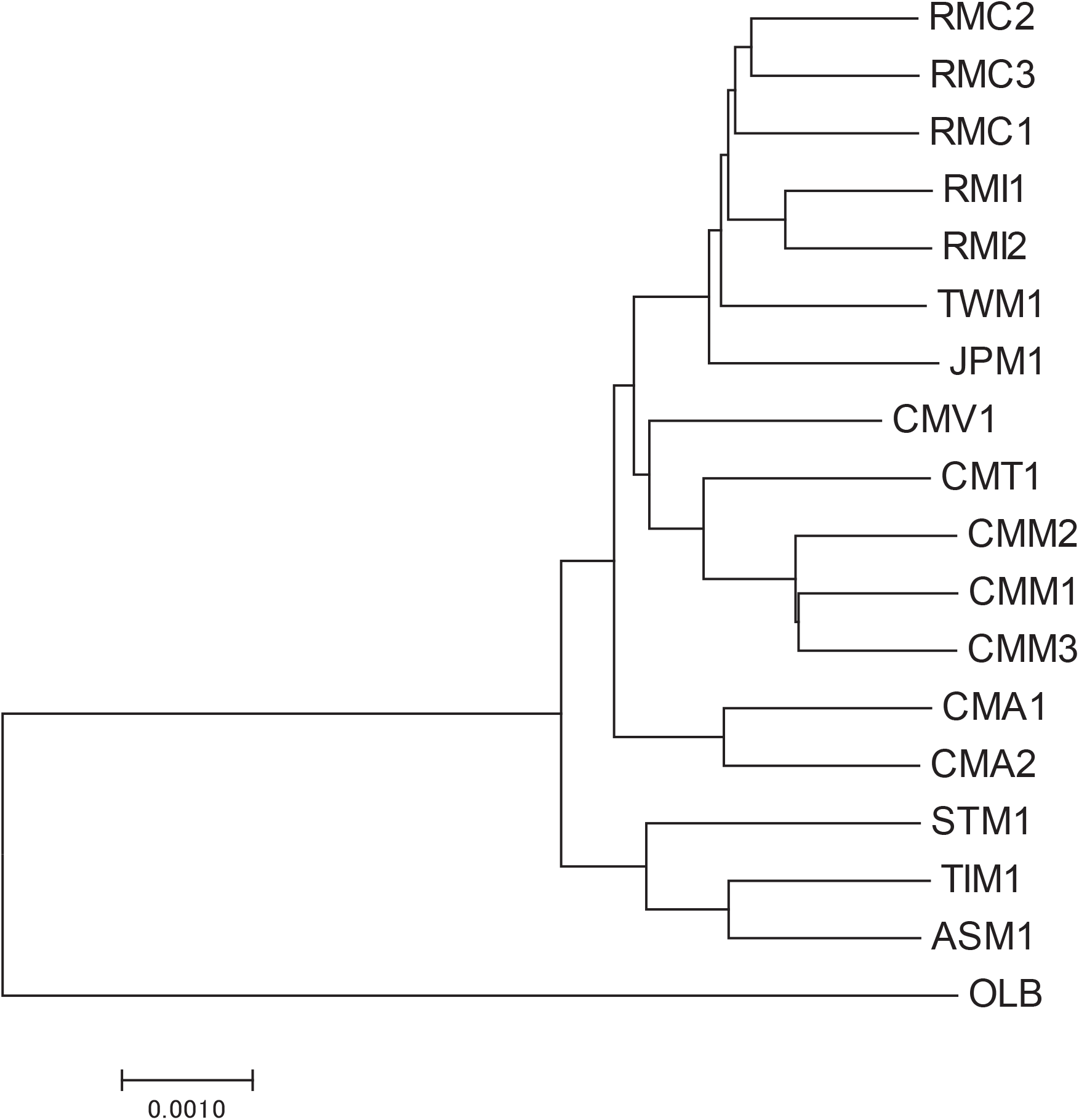
Neighbor-joining tree constructed using the distances between nuclear genomes. All branching patterns are supported by 100 % bootstrap values.

As shown in Figure 3, CMA1 and CMA2 were placed at the root of the *fascicularis*-*mulatta* clades, which is contradicting to the conventional taxonomic classification of *M. fascicularis* ssp. *aurea*. In order to statistically evaluate the genetic relationship between CMA1/CMA2 and other *fascicularis* and *mulatta* group species, we evaluated *f*3 statistics (or outgroup *f*3 statistics) using the OLB genome as the outgroup. In the following analyses based on population genetics theory, we use three-letter symbols to represent the names of the populations to which samples belong. For example, RMC represents the population of which RMC1, RMC2, and RMC3 are members. Although we have only one individual for each of the CMT, CMV, TWM, JPM, STM, TIM, and ASM populations, we can assume that the genotype of the sampled individual reflects the allele frequency of single nucleotide variants (SNVs) in each population. However, we treated samples CMA1 and CMA2 as if they were in different populations, since we do not have any priori assumptions about their population structure. The *f*3 statistics showed that CMA1 and CMA2 were generally closer to CMT and CMV than to CMM or the populations in the *mulatta* group (Figure 4A). For example, the outgroup *f*3 of CMA1 and CMT was significantly greater than that of CMA1 and RMC (*p* = 0.039, *t*-test), supporting that CMA1 and CMA2 were *fascicularis*-like, rather than mulatta-like, populations.

**Figure 4.**
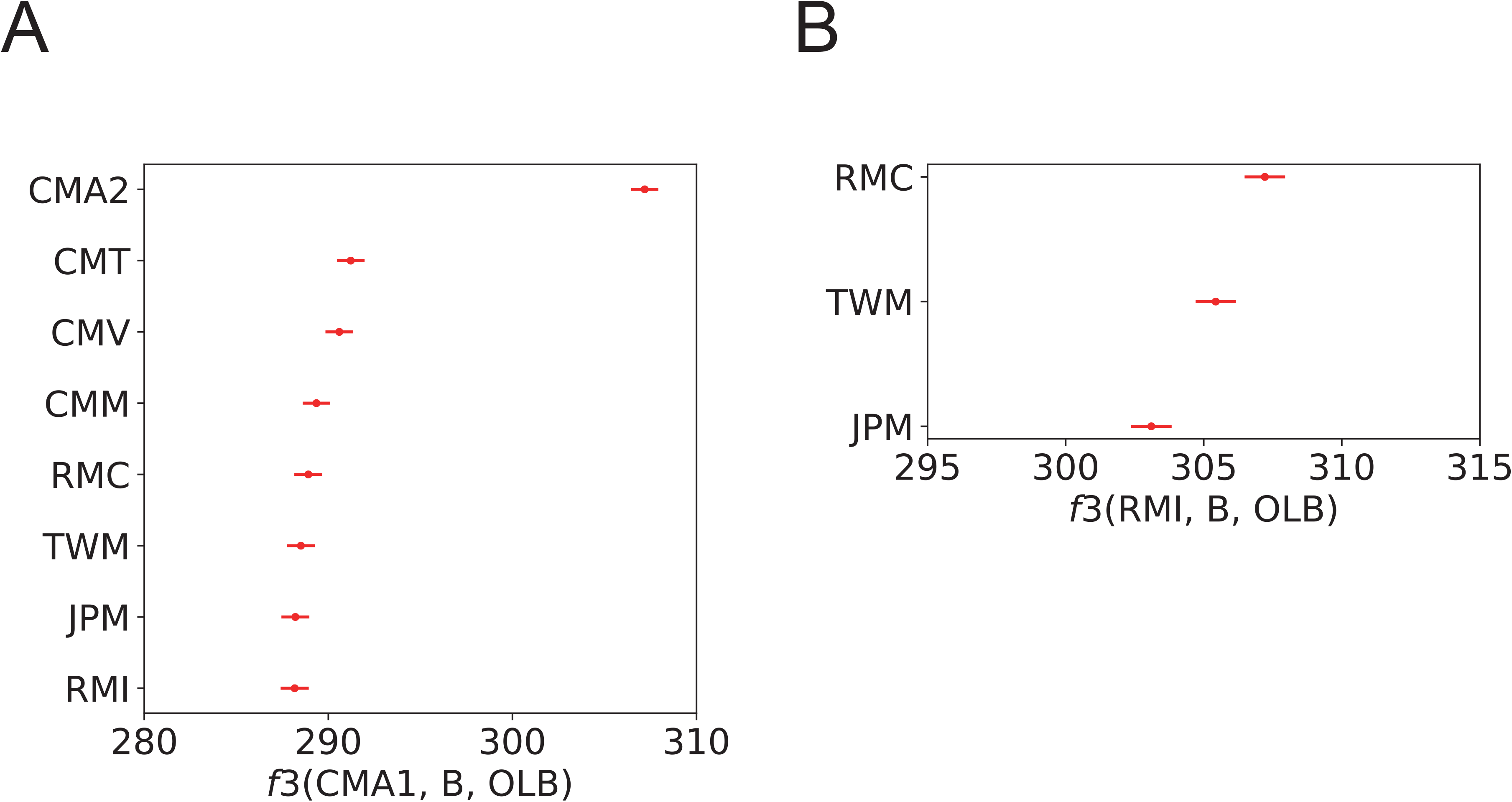
F3 statistics. A) *f*3(CMA1, B, OLB), where population B is shown on the left side of the panel. Large values mean that B is genetically close to CMA1. B) *f*3(RMI, B, OLB), where population B is shown on the left side of the panel. Large values mean that B is genetically close to RMI.

The phylogenetic relationship among the three species of the *mulatta* group—*M. mulatta*, *M. fuscata*, and *M. cyclopis*—has not previously been examined at the whole-genome level. In the neighbor-joining tree shown in Figure 3, the Chinese *M. mulatta* samples clustered with the Indian *M. mulatta* samples, and *M. mulatta* was monophyletic and sister to *M. cyclopis*. However, the branching pattern of the tree may be distorted by gene flow between species/populations. The *f*3 statistics agreed with the pattern in the neighbor-joining tree and indicated that *M. mulatta* and *M. cyclopis* were the sister pair (Figure 4B).

### General picture of admixture between species/populations

We have examined the presence of gene flow between species/populations using *f*4 statistics. Here we represent the configuration of *f*4 statistics as *f*4(A, B; C, D), where A–D represent the names of the populations to which the samples belong. We fixed the species A to OLB, since we assumed that OLB has not experienced any recent gene flow with our macaque populations. As *f*4 statistics are equivalent to *D* statistics, the statistics do not deviate from 0 when the genetic relationship between the four species/populations is represented as a tree structure, indicating no gene flow between species/populations (Patterson et al. 2012). When gene flow occurs between B and D, the *f*4 statistics become positive. When gene flow occurs between B and C, the *f*4 statistics become negative. Hereafter we use the symbol *f*4_A_ to represent *f*4 statistics computed using an autosomal genome.

We computed *f*4_A_(OLB, B; C, D) by choosing B, C, and D for all 220 combinations of sampled populations. Of these, 168 combinations were found to be significant, according to the *f*4 statistics (*p* < 0.0001 after Bonferroni correction) in the best-fitted tree topologies (Supplementary Table 5). These results indicate that the phylogenetic relationships of macaques are highly reticulate, as suggested by Fan et al. (Fan et al. 2018).

The deviation from the tree structure was found to be particularly strong when the configuration included STM, CMA1, CMA2, CMT, and CMV. The all populations in *fascicularis*-*mulatta* group showed a stronger affinity to STM than to the populations in the *sinica* group. For example, *f*4_A_(OLB, RMC; STM, TIM) was strongly negative (*f*4_A_, −0.0019; Z score, −42.9). Similarly, the populations in the *sinica* group were significantly closer to CMA1 and CMA2 than to the populations in the other *fascicularis*-*mulatta* group. Besides, CMV and CMT were more closely related to RMC than to RMI.

### Regional heterogeneity of genetic differentiation on the X chromosomes

Before estimating the level of admixture on the X chromosome, we investigated the regional heterogeneity of genetic differentiation on the X chromosome. We computed *f*4 statistics for each SNV across the non-pseudo-autosomal region of the X chromosome. We found that an approximately 11 Mb length region proximal to the pseudo-autosomal region 1 (PAR1) boundary showed an unusual pattern of differentiation. An example of the *f*4(OLB, ASM; RMI, CMV) across the non-pseudo-autosomal region of the X chromosome is shown in Supplementary Figure 4. The *f*4 values near the PAR1 boundary were observed to be strongly negative and further showed unusually high variance. This pattern was consistently observed in comparisons involving few samples. Although we could not identify the reason for this unusual pattern of differentiation, we excluded the region from further analysis. Hereafter, we designate the *f*4 statistics on the X chromosome excluding the PARs and the 11 Mb region proximal to PAR1 boundary as *f*4_X_.

### Contrasting the patterns of admixture on autosomes and the X chromosome

Comparing the pattern of genetic differentiation between autosomes and X chromosomes, we first examined the level of admixture between STM and the *fascicularis-mulatta*-group populations. The values of *f*4_A_(OLB, B; STM, ASM), where B represents a population in the *fascicularis* or *mulatta* group, were significantly biased in the negative direction (Figure 5A). The *f*4_A_ values of the *mulatta* group were generally lower than those of the *fascicularis* group, indicating a strong affinity of STM to the *mulatta* group.

**Figure 5.**
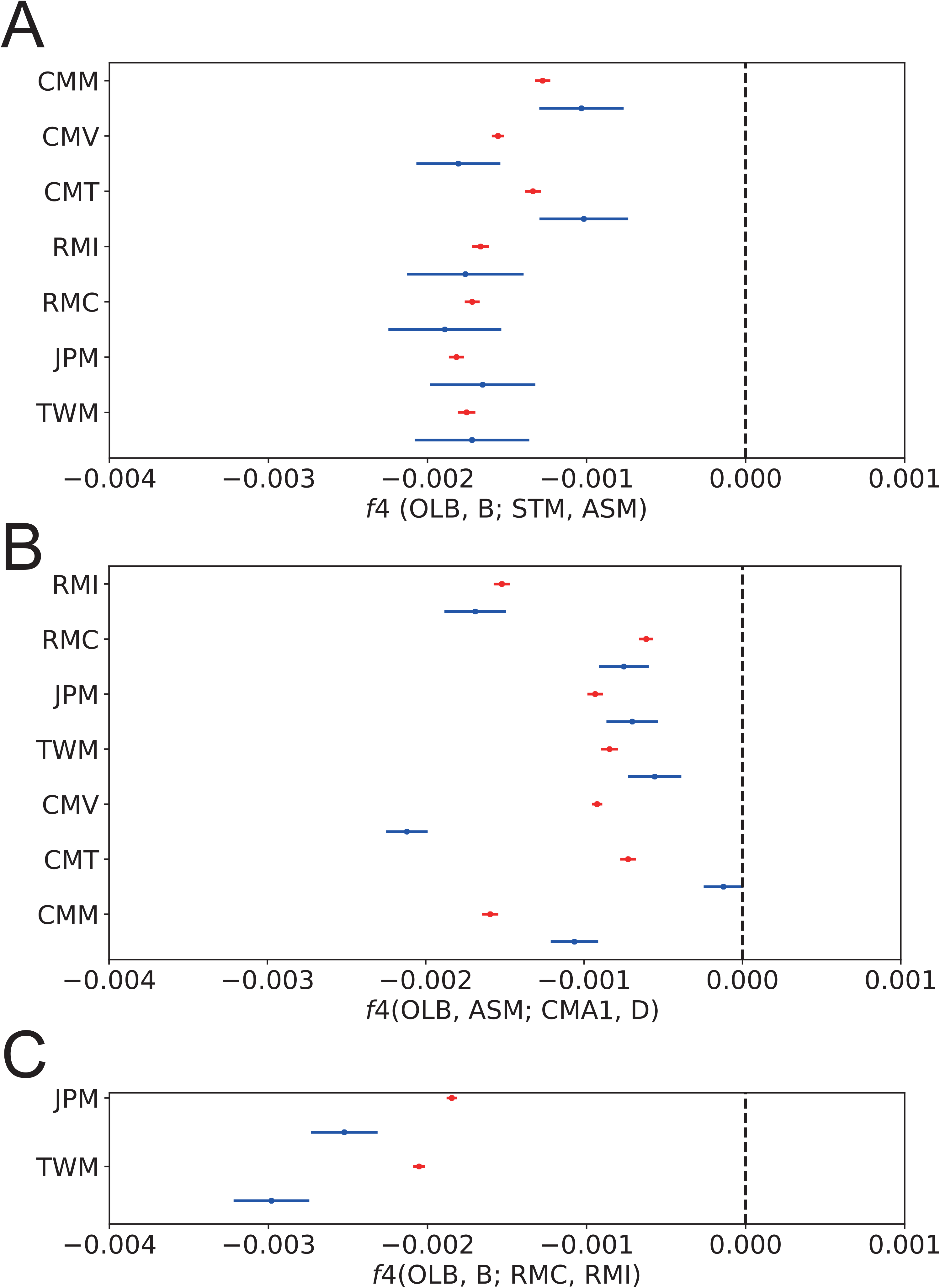
The values of *f*4 statistics on autosomes (red) and the X chromosome (blue). The names of the target species/population are shown on the left side of the panel. A) *f*4(OLB, B; STM, ASM). Negative *f*4 statistics indicate that B is more closely related to STM than to ASM. B) *f*4(OLB, ASM; CMA1, D). Negative *f*4 indicates that ASM is more closely related to CMA1 than to D. C) *f*4(OLB, B; RMC, RMI). Negative *f*4 statistics indicate that B is more closely related to RMC than to RMI.

If the *mulatta*-like mitochondrial genomes of STM originated via nuclear swamping, we could expect that the X chromosomes of STM would be more closely related to those of the *mulatta* group and that *f*4_X_(OLB, B; STM, ASM) would be lower than *f*4_A_(OLB, B; STM, ASM). The rationale for this expectation is presented in Supplementary Figure 5. The results showed that the *f*4_X_ had very large variance across the chromosome and were not significantly different from *f*4_A_. In addition, there was no consistent trend in the differences between the *f*4_X_ and *f*4_A,_ as shown in Figure 5A.

We next compared *f*4_A_ and *f*4_X_ including CMA1 and CMA2. We computed the *f*4(OLB, ASM; CMA1/2, D), where D represents a population in the *fascicularis* or *mulatta* group other than CMA1 and CMA2. The *f*4_A_ values were observed to be strongly negative, indicating that ASM was significantly more closely related to CMA1 and CMA2 than to the other *fascicularis-* and *mulatta*-group populations (Figure 5B), which coincides with the observation that the mitochondrial genomes of CMA1 and CMA2 clustered with those of the *sinica* group (Figure 2).

Similar to the previous analysis, if the *sinica*-like mitochondrial genomes of CMA1 and CMA2 were the consequence of nuclear swamping, we would expect that the X chromosomes of CMA1 and CMA2 would be more closely related to those of the *sinica* group than to the autosomes, resulting in *f*4_X_ being lower than *f*4_A_. In one comparison, *f*4_X_(OLB, ASM; CMA1, CMV) was significantly lower than *f*4_A_(OLB, ASM; CMA1, CMV), supporting the nuclear swamping hypothesis; in the other comparisons, however, the *f*4_X_ values were not significantly different from or were significantly higher than those on autosomes. Again, the results did not consistently support the nuclear swamping hypothesis.

We finally looked at the *f*4 statistics to detect admixture among the species/populations in the *mulatta* group. We computed *f*4_A_ and *f*4_X_ values for *f*4(OLB, JPM; RMC, RMI) and *f*4(OLB, TWM; RMC, RMI). In both cases, the *f*4_A_ and *f*4_X_ values were significantly negative, indicating that JPM and TWM were more closely related to RMC than to RMI (Figure 5C). Since *M. fuscata* and *M. cyclopis* do not form a sister species pair (Figure 3), this result indicates that there were at least two rounds of past admixture events among the ancestral populations of Chinese *M. mulatta*, *M. fuscata*, and *M. cyclopis*. This pattern of admixture is consistent with the mitochondrial phylogeny, in which individuals of Chinese *M. mulatta* clustered with JPM1 and TWM1. We also found that *f*4_X_ was significantly lower than *f*4_A_ (*p* = 0.0044 for JPM and *p* = 0.0008 for TWM, *t*-test), which shows that RMC were more similar to JPM and TWM on the X chromosomes than on autosomes, a pattern expected under the nuclear swamping hypothesis

To support these observations, we constructed an admixture graph using autosomal data. The admixture graph suggested a complex admixture history of the *mulatta* group (Figure 6). Although the reconstructed admixture graph is only one of the good past demographic models that explains the data, the admixture graph was concordant with the hypothesis of two-round hybridization for the origin of Chinese *M. mulatta* inferred by the *f*4 statistics. Based on this model, the populations that diverged from the lineages of *M. fuscata* and *M. cyclopis* generated a hybrid population (node K in Fig. 6), and later, this hybrid population admixed with the ancestral population of Chinese *M. mulatta* (node O in Fig. 6).

**Figure 6.**
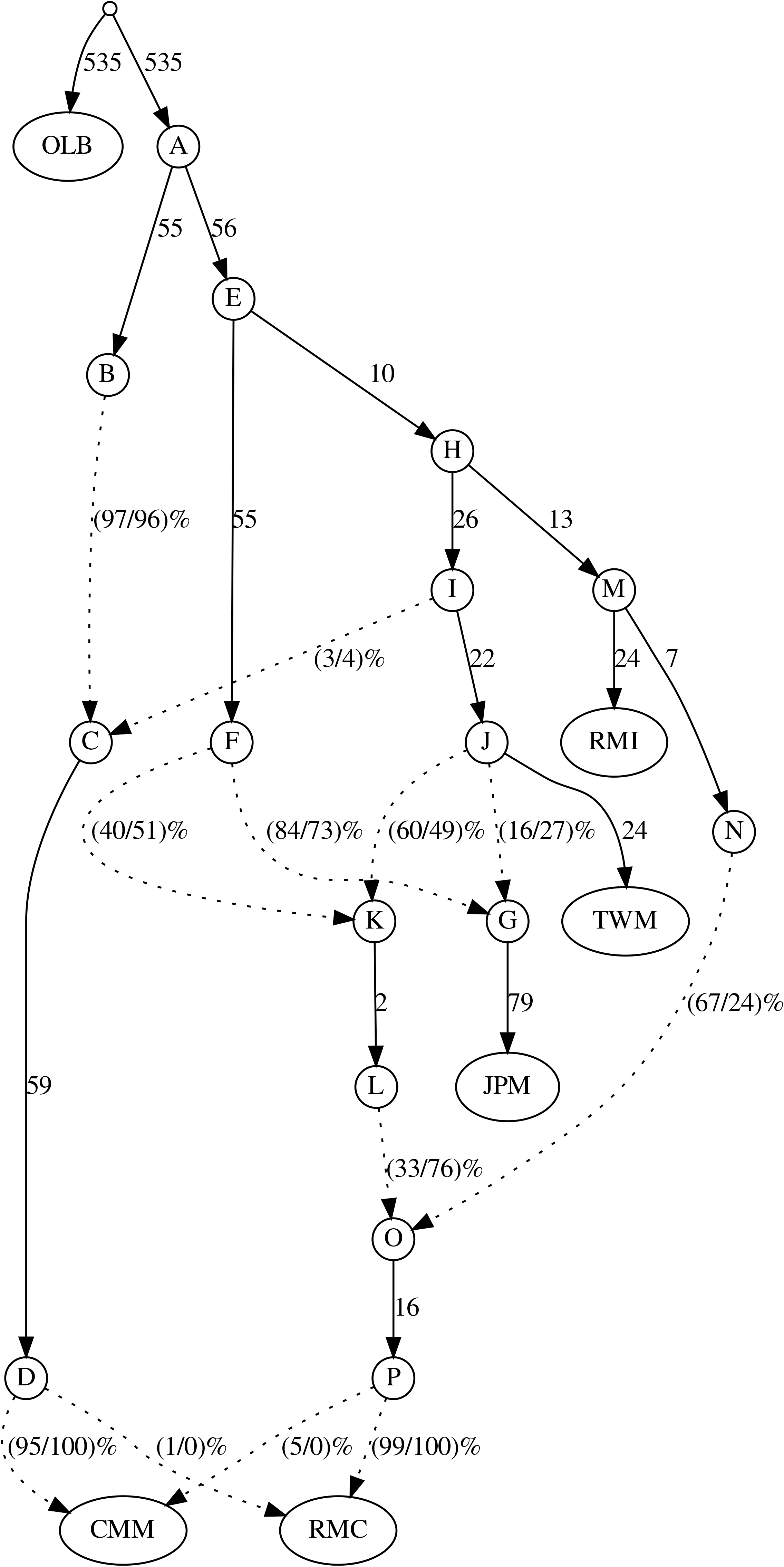
Admixture graph of the *mulatta*-group species and CMM. OLB was used as the outgroup. The numbers along the branches (solid lines) indicate drift parameters estimated using autosomes (left) and X chromosomes (right). The dashed lines represent admixture events, and the numbers in parenthesis show the admixed fraction (%) estimated using autosomes (left) and X chromosomes (right). In this figure, the establishment of the RMC population is explained by two admixture events. (1) The admixture of populations derived from the JPM lineage (node F) and a population derived from TWM lineage (node J) created a hybrid population (node K). (2) The admixture of the hybrid lineage (node L) and a population diverged from RMI (node N) generated the ancestral population of RMC. Note that the admixture fraction of node N is much larger on autosomes than on X chromosomes (67 % and 24 % on autosomes and X chromosomes, respectively). This admiture graph did not show any statistically significant deviation.

We assume that the admixture graph inferred from the autosomal data reflects the past demographic events among species reasonably accurately and estimated the admixed fraction on the X chromosome by fixing the admixture graph (Figure 6). In general, the level of admixture on the X chromosomes was lower than that on autosomes. However, the admixed fraction from the ghost population (node L in Fig. 6) in the RMC genomes on the X chromosomes was 67 %, which was much higher than that on autosomes (33%). This result is consistent with the pattern observed in the analysis of the *f*4 statistics, and this also supports the nuclear swamping hypothesis, which posits that strong, continuous male-biased migration from the ancestral Chinese *M. mulatta* population (node N in Fig. 6) to the ghost population diverged from *M. fuscata* and *M. cyclopis* (node L in Fig. 6) generated incongruencies between the genealogies of the mitochondrial and autosomal genomes.

## Discussion

In this study, we examined 17 macaque genomes to investigate their evolutionary history. Comparing the pattern of genetic differentiation between the autosomal and X-chromosomal genomes is the primary focus of this study, in order to quantify the effect of sex-biased migration, which is universally observed in sexually reproducing organisms. The number of samples per species in this study was limited, and we mostly neglected the genetic structure within species. In particular, *M. mulatta* and *M. fascicularis* are widely distributed, and they showed detectable levels of genetic structure within the species (Bunlungsup et al. 2017; Liu et al. 2018). Therefore, a more detailed and comprehensive evolutionary history of macaques should be performed using species-wide genome sequences in future studies. Despite this limitation, our analysis including five newly sequenced genomes revealed many novel aspects of macaque evolutionary history.

The genetic relationship of *M. fascicularis* ssp. *aurea* to other species has only been recently studied using mitochondrial and Y-chromosomal genomes (Matsudaira et al. 2018). It was reported that *M. fascicularis* ssp. *aurea* used stone tools for foraging (Carpenter 1887; Malaivijitnond et al. 2007), a behavior which is unique to the subspecies. Our genome-wide analysis suggested a highly complex scenario for the origin of the subspecies. Strong admixture was detected between *M. fascicularis* ssp. *aurea* and the *sinica* group; this implies that *M. fascicularis* ssp. *aurea* originated from the ancient hybridization with a population related to *sinica*-group species. *f*3 statistics indicated that CMA1 and CMA2 are most closely related to CMT and CMV (Figure 4A). In contrast, the genetic distance between CMA1/2 and CMM was not significantly different from that between CMA1/2 and *mulatta*-group populations. It is difficult to determine whether the *M. fascicularis* ssp. *aurea* lineage branched after or before the split of the *fascicularis* and *mulatta* groups because of substantial gene flow between species. Our results showed that the genomic features of *M. fascicularis* ssp. *aurea* are unique among the *M. fascicularis* samples. CMA1 had very low genetic diversity compared with the other *M. fascicularis* individuals in this study. The large number of ROH in CMA1 implies that the reduction of genetic diversity may be due to recent inbreeding. Indeed, CMA1 was sampled from a population living on a small island, and the population size remained the same for decades.

The genome-wide phylogenetic relationship among species in the *mulatta* group was also clarified in this study. Using approximately 8 Mb of autosomal sequence data, Perelman et al. (Perelman et al. 2011) reported a sister relationship between *M. mulatta* and *M. fuscata*. However, our analysis showed that *M. mulatta* is closer to *M. cyclopis* than to *M. fuscata* at autosomal loci (Figures 3 and 5C), a finding which is concordant with those of a study using isozymes by Melnick et al. (Melnick and Kidd 1985). In addition, *f*4 statistics showed a high level of admixture between Chinese *M. mulatta* and *M. fuscata*/*cyclopis*. Since *M. fuscata* and *M. cyclopis* were found to not form a sister pair, there have been multiple admixture events among the *mulatta*-group species. In order to provide deeper insights into the evolutionary history of the *mulatta*-group species, we re-analyzed population genomics data produced by Liu et al. (2018) from the Chinese *M. mulatta*, including five subspecies: *M. m. tcheliensis*, *M. m. littoralis*, *M. m. brevicaudus*, *M. m. lasiotis*, and *M. m. mulatta*. One individual of each subspecies was added to our dataset (Supplementary Table 6). Although all the subspecies showed strong signatures of admixture with *M. fuscata* and *M. cyclopis*, significant heterogeneity in the relatedness of Chinese *M. mulatta* to *M. cyclopis* and *M. fuscata* was found; *f*4 statistics showed that the subspecies *M. m. tcheliensis* from North China and *M. m. mulatta* from Yunnan were more dissimilar to *M. cyclopis* and *M. fuscata* than the other subspecies (Supplementary Figure 6). The two subspecies currently inhabit peripheral regions to the range of *M. mulatta*. Since *M. m. tcheliensis* would be the latest derived subspecies (Liu et al. 2018), the process of admixture would have continued in multiple regions around Southern China.

In an evolutionary history proposed by Delson (Delson 1980), the most recent common ancestors of *M. mulatta* lived around Eastern India or Myanmar and split to form the Indian and Chinese populations. Using genome-wide polymorphism data, Hernandez et al. estimated the split to occur around the Middle to Late Pleistocene boundary (162 kya) (Hernandez et al. 2007). After the split, the Chinese population expanded eastward and northward as their population size increased. Both mitochondrial and X-chromosomal patterns were explained by the hypothesis that strong male-biased migration from the ancestral Chinese *M. mulatta* population toward the ghost population related to *M. fuscata* and *M. cyclopis* occurred after the split between the Chinese and Indian populations. This scenario is also supported by the reconstructed admixture graph (Figure 6). We show the hypothetical evolutionary history of the *mulatta*-group species in Figure 7.

**Figure 7.**
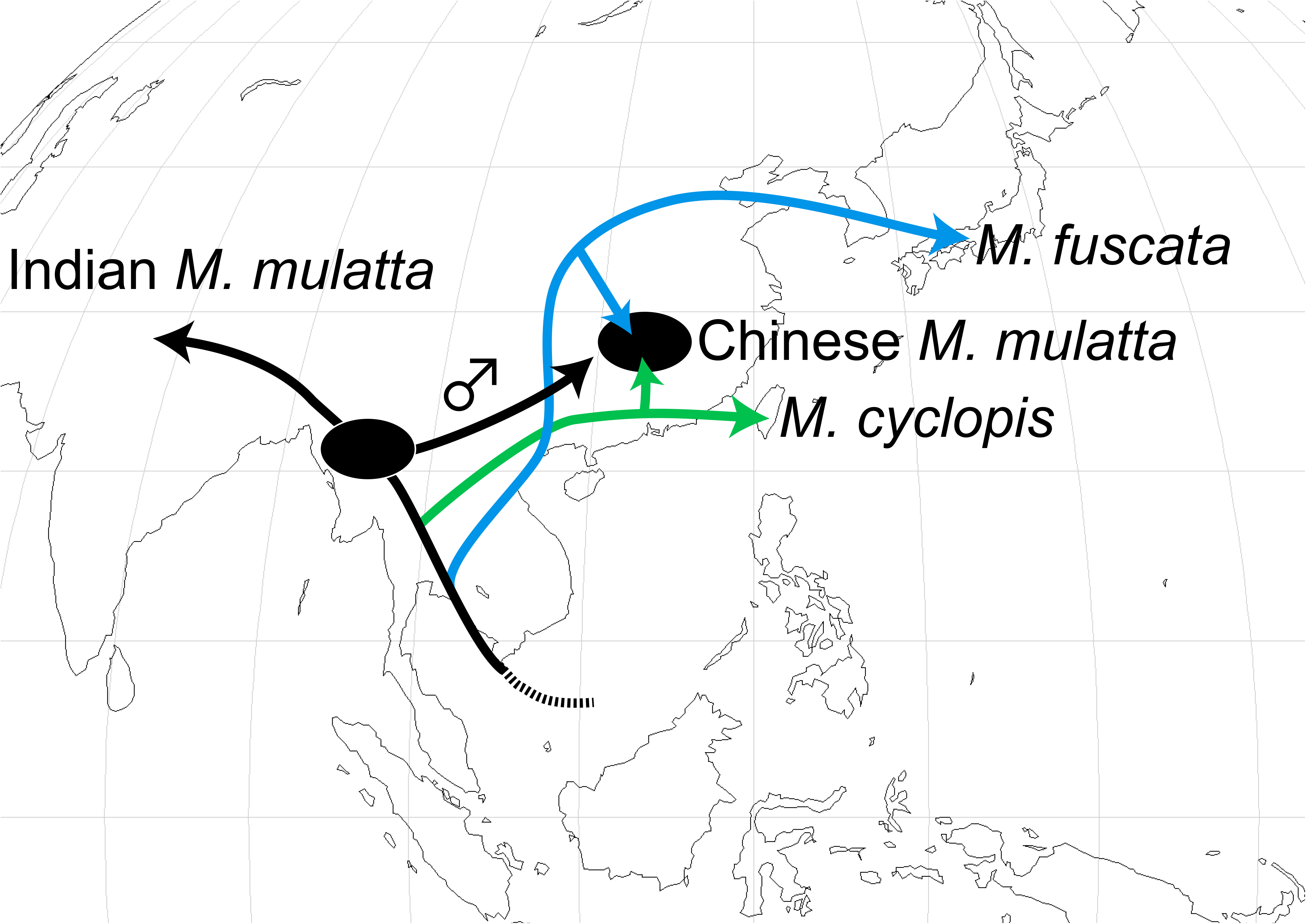
A hypothetical evolutionary history of the *mulatta*-group species. Arrows indicate the direction of migration. We propose the existence of a ghost population generated by the admixture of populations diverged from the *M. fuscata* and *M. cyclopis* lineages. Strong male-biased migration from an ancestral Chinese *M. mulatta* population (represented by the symbol ♂) to the ghost population established the genetic features of extant Chinese *M. mulatta*. The gene flow between Chinese and Indian *M. mulatta* was detected in the previous study (Hernandez et al. 2007) but is omitted in this figure.

There are significant levels of gene flow among these species. Of the 220 possible configurations of *f*4 statistics, 169 have showed deviation from a simple phylogenetic tree (*p* < 0.0001). The reason for the large number of deviant comparisons might be that the *f*4 test is too sensitive. We tested the deviation of *f*4_A_(OLB, B; C, D), where C and D represent individuals from the same populations, and estimated the degree of deviation from treeness. We found that some comparisons within the population showed a statistically significant deviation from the expectation. For example, *f*4_A_(OLB, STM; CMM1, CMM2) was −2.1×10^−4^ (Z score, −11.9), which was highest deviation among the comparisons. We cannot confirm whether these deviations are due to technical artifacts or to hidden population structure. However, most of the observed degrees of deviation in our main analysis were much higher than these values; 100 out of the 220 comparisons had a Z score greater than 20 or smaller than −20, indicating that most of the inferred admixture events accurately reflect past demographic events.

For *M. arctoides* and *M. fascicularis* ssp. *aurea*, we did not find evidence supporting the nuclear swamping hypothesis for their origination. Given the significant gene flow among species, the evolutionary history of the *fascicularis* and *sinica* groups would be extremely complicated and may have blurred the signature of the past sex-biased migration. Alternatively, the incongruence between autosomal and mitochondrial genealogies could be explained by the introgression of mitochondrial genomes. Studies with larger sample sizes may be conducted to answer the question in the near future.

We should also note that the different patterns of genetic differentiation between autosomal and X-chromosomal loci expected under nuclear swamping are also explained by strong natural selection against the introgression of X chromosomes. These factors are not mutually exclusive, and their relative importance should be examined using multiple sources of evidence. In the case of Chinese *M. mulatta*, if we assume that the mitochondrial locus is evolutionarily neutral, the pattern observed provides evidence supporting the nuclear swamping hypothesis. However, several studies have suggested that the introgression of mitochondrial genomes might be deleterious because they may disrupt the interactions between nuclear and mitochondrial alleles, a phenomenon which is referred to as mitonuclear incompatibility (Osada and Akashi 2012; Hill 2020). In such a scenario, the patterns expected under male-biased migration and selection against X-chromosomal and mitochondrial introgression would be indistinguishable. Therefore, in order to understand the factors shaping the diversity of genomes and elucidate the natural history of organisms, population genomic studies on a wide range of organisms with different sex-determination systems and different migration patterns are needed.

## Conclusions

In this study, we investigated the complex evolutionary history of macaques with rampant gene flow among species, using a whole-genome sequence dataset. As shown in previous studies, incongruencies in genealogical relationships between nuclear and mitochondrial genomes were observed among species. We showed that comparing the pattern of admixture between autosomal and X-chromosomal loci is a potentially valuable approach to statistically evaluate the power of sex-biased migration in shaping the pattern of genome evolution. We detected a statistically significant difference in admixture levels between these loci, which could be explained by strong male-biased migration in one of the three cases tested. Comparisons between an autosomal and an X-chromosomal evolutionary pattern using a larger dataset in other species would reveal a more detailed evolutionary history in future studies.

## Materials and Methods

### Sample collection and genome sequencing

DNA samples of *M. fuscata* and *M. cyclopis* were obtained from the Primate Research Institute (PRI), Kyoto University. The monkeys were cared for and handled according to the guidelines established by the Institutional Animal Welfare and Animal Care Committee of PRI. They were anesthetized, and peripheral blood was obtained. The DNA samples were used for library preparation, and paired-end sequences of 101 bp were determined using HiSeq 2500 (Illumina, CA, USA). All experimental procedures were approved by the Institutional Animal Welfare and Animal Care Committee of PRI (No. 2015-138).

The three *M. fascicularis* samples were obtained from temporally-caught wild animals in Thailand. The animals were anesthetized, and blood samples were withdrawn from the femoral vein. The protocol was approved by the Institutional Animal Care and Use Committee of the Faculty of Science, in accordance with the guidelines for the care and use of laboratory animals prepared by Chulalongkorn University, Protocol Review No. 1423010. DNA was extracted from the buffy coat using a standard phenol-chloroform method as described in a previous study (Bunlungsup et al. 2016). The native DNA was whole-genome amplified (WGA) and substituted to artificially synthesized DNA. WGA was conducted using the REPLI-g Mini Kit (Qiagen, Hilden, Germany) following the manufacturer’s protocol. The WGA products were purified using Wizard® SV Gels and PCR Clean-Up System (Promega, WI, USA). The DNA samples were used for library preparation, and paired-end sequences of 151 bp were determined using HiSeq X (Illumina, CA, USA).

All short-read sequences were deposited in the public database DDBJ/DRA under the project ID PRJDB9555. Detailed sample information is presented in Supplementary Table 1.

### Variant calling

For all samples, reads were mapped to the reference genome sequence of OLB (Panu_3.0; accession number: GCA_00264685.2) using the BWA-MEM algorithm (Li and Durbin 2009) with default parameter settings. Short reads potentially derived from PCR duplication were labeled using Picard software. SNVs were called using HaplotypeCaller in GATK 4.0 with a prior probability of heterozygosity of 0.003 and a cut-off quality score of 30. The SNVs were further hard-filtered with the following parameters: QD < 2.0, FS > 60.0, SOR > 9.0, MQ < 40.0, MQRankSum < −12.5, and ReadPosRankSum < −8.0. For SNV calling on X chromosomes, males and females were genotyped separately, according to the ploidy of the X chromosome. The heterozygosity of the samples was calculated using an in-house-generated Python script, which calls the number of homozygous and heterozygous nucleotide sites with coverage between 10 and 60. The segments of the ROH were estimated using the PLINK software (Purcell et al. 2007).

### Phylogenetic tree of mitochondrial genomes

We assembled mitochondrial genomes from short-read sequences using the NOVOPlasty software (Dierckxsens, Mardulyn, and Smits 2016). Of the 17 samples we analyzed, 9 yielded circular mitochondrial genome assemblies. The five newly determined mitochondrial genome sequences were deposited in the public databases (DDBJ/EMBL/NCBI accession numbers: LC536644–LC536648). The mitochondrial genomes of CMV1 was assembled using the same method with the data obtained by Yan et al. (Yan et al. 2011). The mitochondrial genomes of *M. nigra*, Bornean *M. nemestrina*, and *M. tonkeana* were assembled using the same method, with the data obtained by Evans et al. (Evans et al. 2017). Thirty-one whole mitochondrial genomes of other macaque species and that of *P. anubis* were downloaded from the public database. The DDBJ/EMBL/GenBank accession numbers of sequences are given in Supplementary Table 2. A phylogenetic tree was reconstructed using MEGA X with the maximum likelihood method (Knyaz et al. 2018). The HKY substitution and G+I (gamma + invariant sites) rate models were used.

### Analysis of species phylogeny and admixture

To construct a neighbor-joining tree and neighbor-net network, all pairwise genomic distances were calculated. The genomic distance *D* between two diploid genomes, X and Y, was calculated using the following equation.

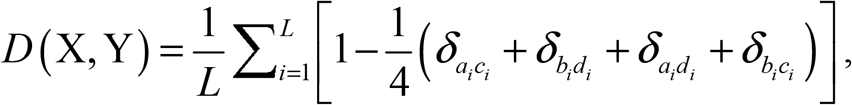

where *a_i_b_i_* and *c_i_d_i_* represent the genotypes of X and Y at the *i*-th site and *δ_pq_* indicates Kronecker delta. The neighbor-net network was constructed using the phangorn package in R (Schliep 2010).

Population trees with migration edges were reconstructed using the TreeMix software with a block size parameter (k) of 500 (Pickrell and Pritchard 2012). f3 and f4 statistics were calculated using the AdmixTools software package (Patterson et al. 2012). Admixture graphs were constructed using the qpGraph software in the AdmixTools software package.

## Acknowledgments

We are grateful to Srichan Bunlungsup and Zhenxin Fan for collaboration on the early stages of this work. We would also like to thank Enago (www.enago.jp) for the English language review. This work was supported by the Japan Society for the Promotion of Science KAKENHI (grant number 18H05511 to N.O.), the Thailand Research Fund-Chinese Academy of Science (grant number DBG60 to S.M.), and Thailand Research Fund Senior Scholar (grant number RTA6280010 to S.M.).

## Notes

### Competing Interest Statement

The authors have declared no competing interest.

