## Supplementary Figures for "Sex-biased migration and admixture in macaque species revealed by comparison between autosomal and X-chromosomal genomic sequences"

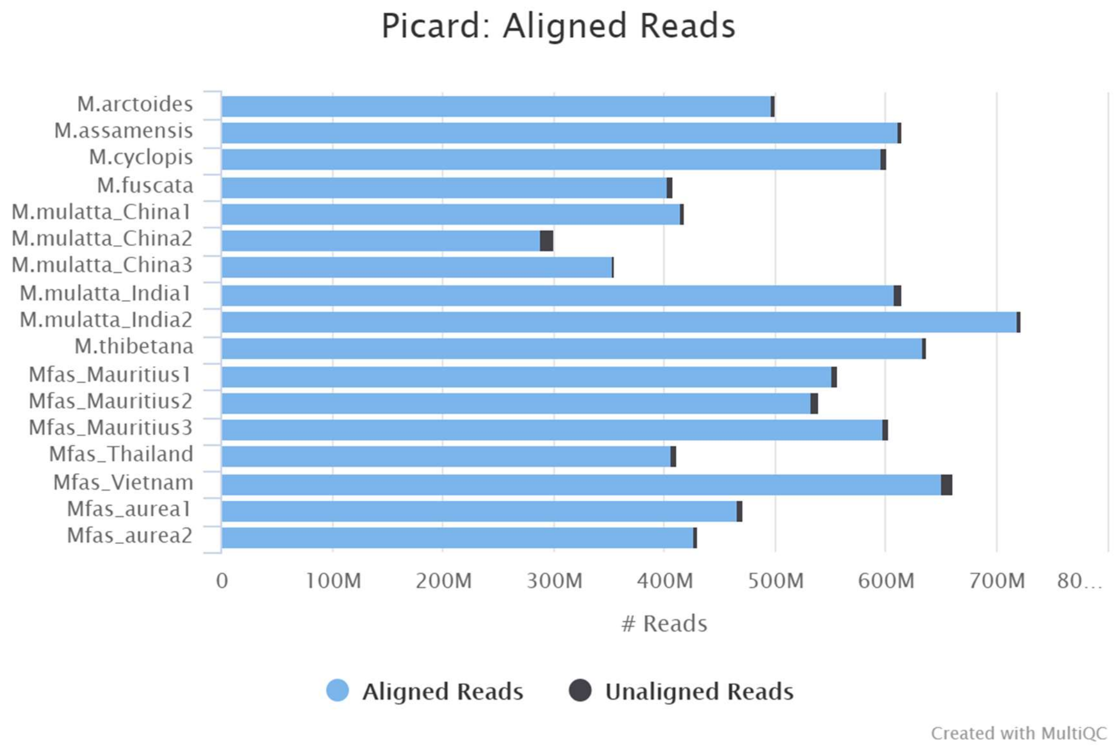


**Supplementary Figure 1**

Numbers of the total and mapped reads, generated using the MultiQC software (Ewels et al. 2016).

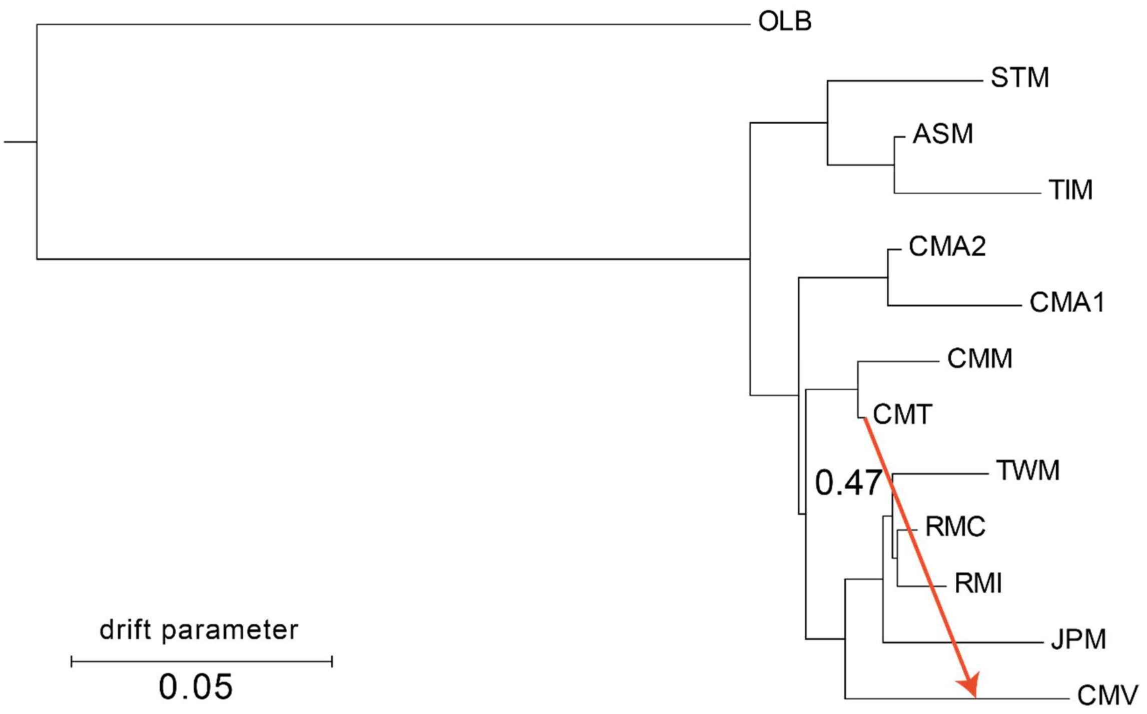


**Supplementary Figure 2**

Population tree reconstructed using TreeMix. One migration edge was added.

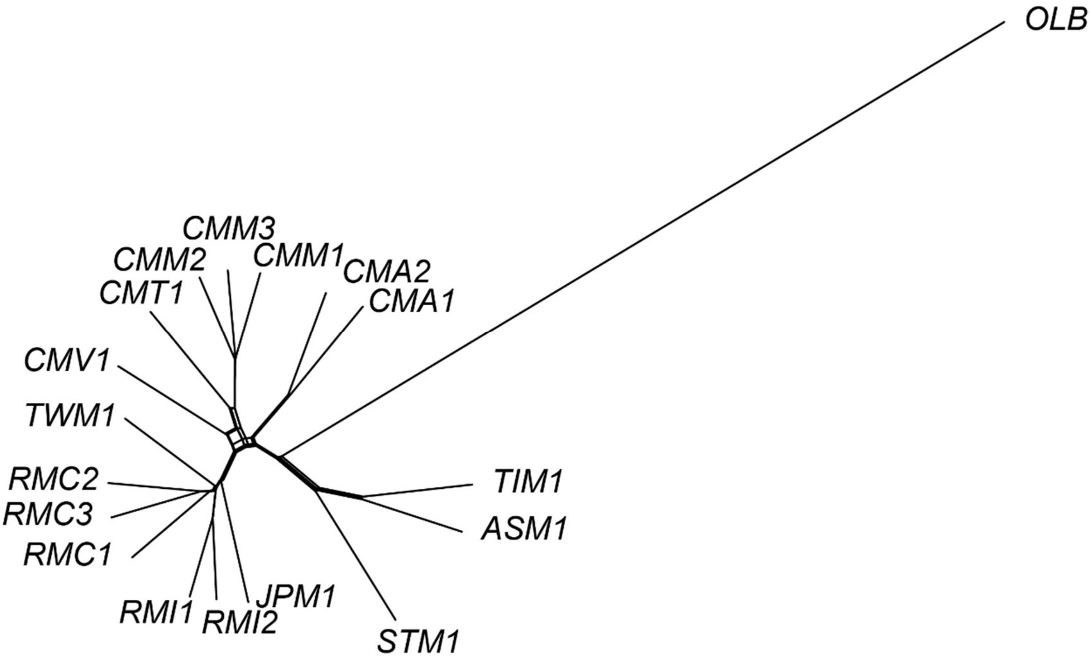


**Supplementary Figure 3**

Neighbor-net network. The names of samples are shown in Table 1 in the main text.

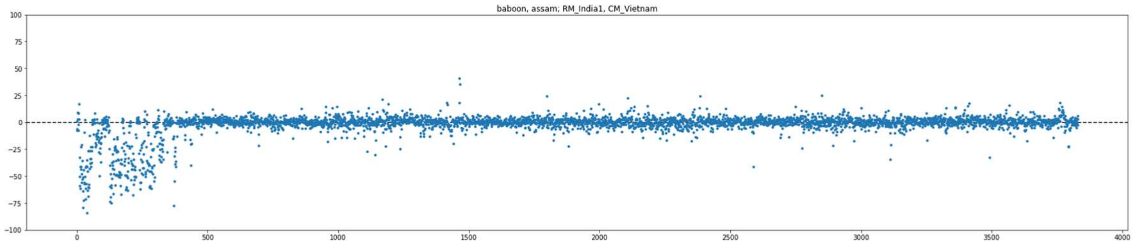


**Supplementary Figure 4**

Distribution of $f4_X$(OLB, ASM; RMI, CMV) across the non-pseudo-autosomal region of the X chromosome. The *x*- and *y*-axes represent the order of SNV sorted by the genome coordinates and *f*4 values, respectively.

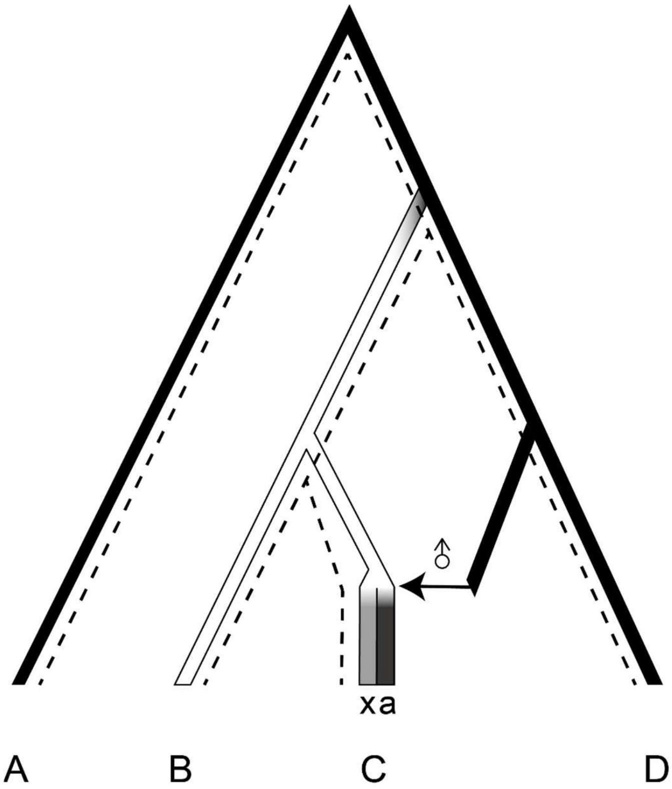


$$0 > f4_A(A, B; C, D) > f4_X(A, B; C, D)$$

**Supplementary Figure 5**

Schematic illustration of testing the nuclear swamping model using *f*4 statistics. Here, we analyze four populations, A–D, where A is the outgroup population. The tick bars represent the history of nuclear genomes, and the dashed lines show mitochondrial genealogy. In this scenario, the common ancestors of B and C split from the lineage of D, and genetic drift has changed allele frequencies in populations (represented by black and white colors). After the divergence of B and C, we assume strong and continuous male-biased migration events from the sister population of D (the horizontal arrow). Finally, the autosomal and Y-chromosomal genome was almost replaced by the one from the donor population (the dark gray bar over "a"). On the other hand, the replacement of X-chromosomal genome would be weak so that the X chromosomes retained original genetic variation (the light gray bar over "x"). When the replacement is not complete, $f4_A$(A, B; C, D) becomes negative, but the deviation would be milder than $f4_X$(A, B; C, D).

A

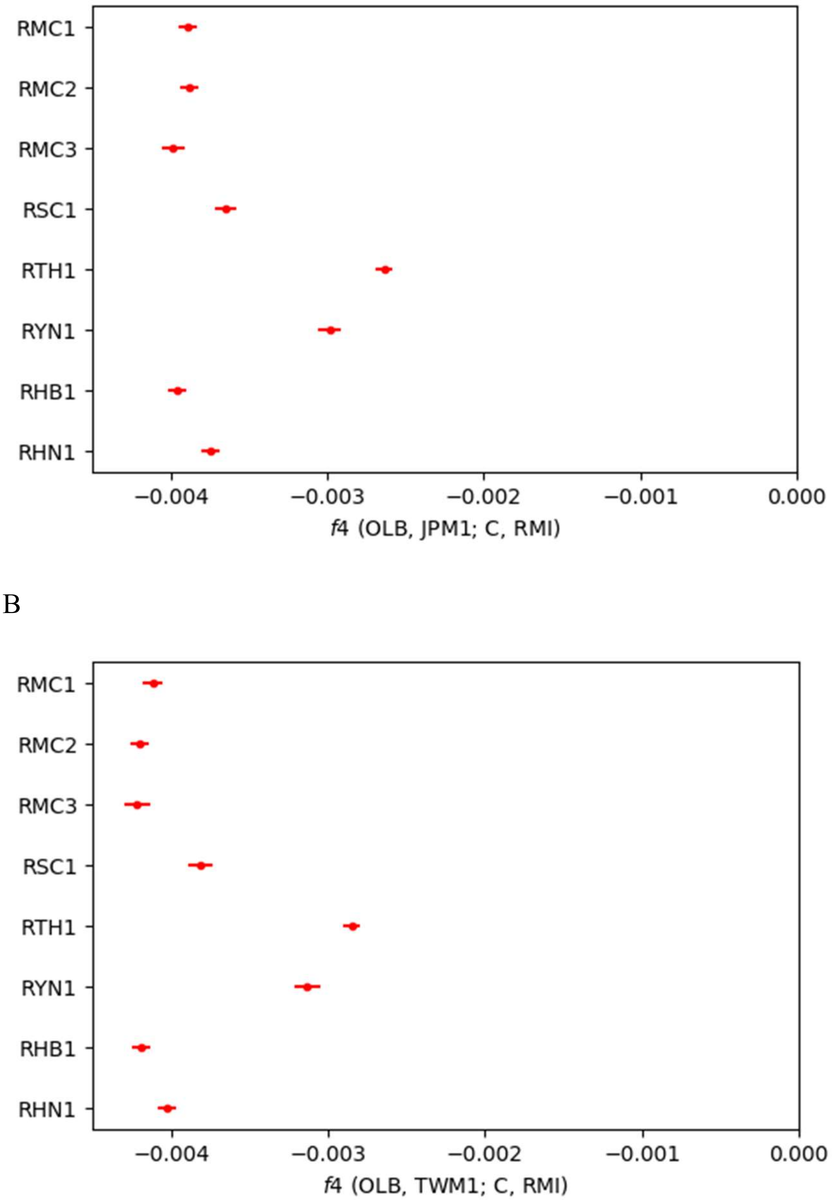


**Supplementary Figure 6**

The values of *f*4 statistics on autosomes. The names of the target species/population are shown on the left side of the panel. A) *f*4(OLB, JPM; C, RMI). Negative *f*4 statistics represent that JPM is more closely related to C than RMI. B) *f*4(OLB, TWM; C, RMI). Negative *f*4 represents that TWM is more closely related to C than RMI. RSC, *M. m. lasiotis*; RTH, *M. m. tcheliensis*; RYN, *M. m. mulatta*; RHB, *M. m. littoralis*, RHN, *M. m. brevicaudus*.
